# Luminal Flow Actuation Generates Coupled Shear and Strain in a Microvessel-on-Chip

**DOI:** 10.1101/2021.04.10.439271

**Authors:** Claire A. Dessalles, Clara Ramón-Lozano, Avin Babataheri, Abdul I. Barakat

## Abstract

In the microvasculature, blood flow-derived forces are key regulators of vascular structure and function. Consequently, the development of hydrogel-based microvessel-on-chip systems that strive to mimic the in vivo cellular organization and mechanical environment has received great attention in recent years. However, despite intensive efforts, current microvessel-on-chip systems suffer from several limitations, most notably failure to produce physiologically relevant wall strain levels. In this study, a novel microvessel-on-chip based on the templating technique and using luminal flow actuation to generate physiologically relevant levels of wall shear stress and circumferential stretch is presented. Normal forces induced by the luminal pressure compress the surrounding soft collagen hydrogel, dilate the channel, and create large circumferential strain. The fluid pressure gradient in the system drives flow forward and generates realistic pulsatile wall shear stresses. Rigorous characterization of the system reveals the crucial role played by the poroelastic behavior of the hydrogel in determining the magnitudes of the wall shear stress and strain. The experimental measurements are combined with an analytical model of flow in both the lumen and the porous hydrogel to provide an exceptionally versatile user manual for an application-based choice of parameters in microvessels-on-chip. This unique strategy of flow actuation adds a dimension to the capabilities of microvessel-on-chip systems and provides a more general framework for improving hydrogel-based in vitro engineered platforms.

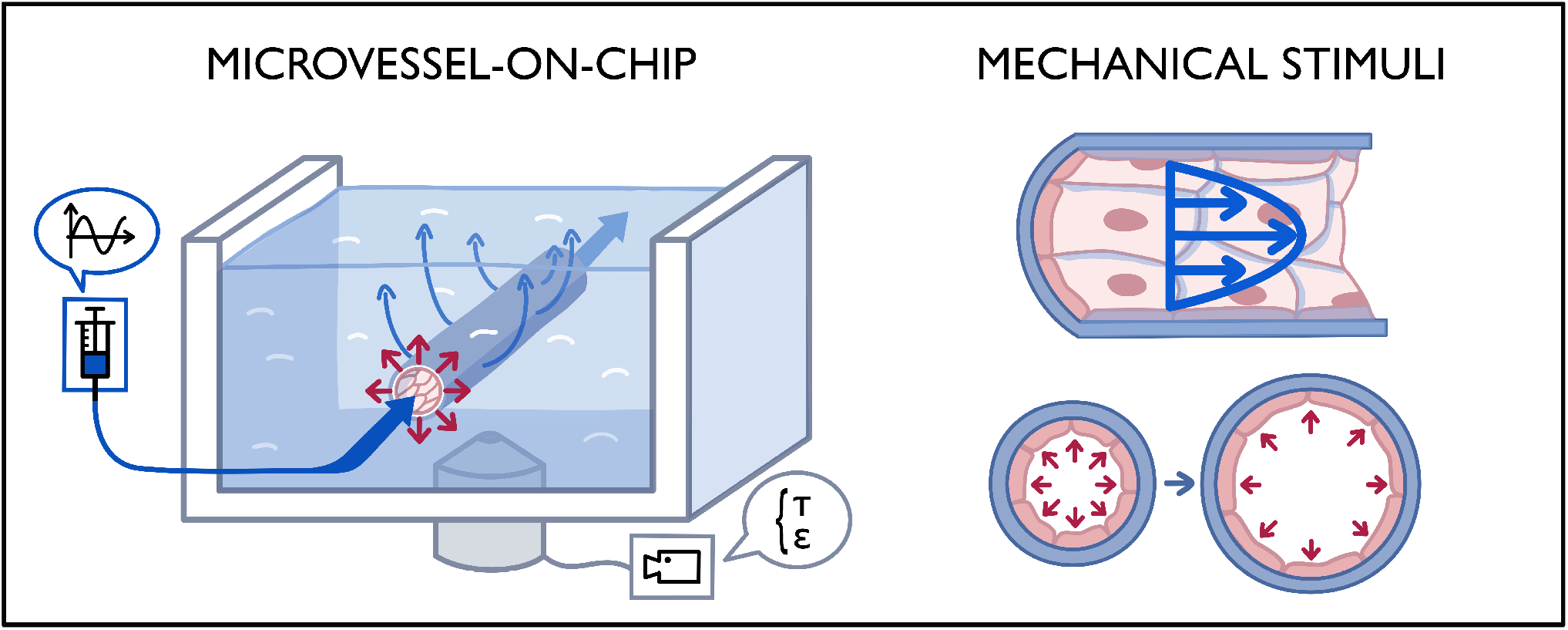

## Introduction

In the vasculature, mechanical forces regulate blood vessel development, network architecture, wall remodeling, and arterio-venous specification [1, 2]. Abnormalities in the mechanical environment play a critical role in the development of vascular diseases in both large and small vessels. In the specific case of the microvasculature, hemodynamic perturbations are associated with chronic pathologies such as hypertension as well as more acute events such as thrombosis [2–5]. Vascular responsiveness to mechanical stimulation is enabled in large part by the endothelium, the layer of cells at the interface between the bloodstream and the vascular wall. By virtue of their strategic position, endothelial cells (ECs) sense mechanical forces and transduce these forces into signals that regulate vascular structure and function [6–8].

Elucidating the mechanisms governing endothelial mechanobiology requires the development of in vitro systems within which ECs experience a physiologically relevant mechanical environment [9, 10]. In vivo, the endothelium is principally subjected to two types of mechanical stresses that act in different directions: tangential shear stress due to the flow of viscous blood over the EC luminal surface [11] and circumferential (hoop) stress as a result of the transmural pressure difference which generates a circumferential strain that stretches the cells. Typical physiological shear stress values range from 0.5 to 5 Pa, depending on the location in the vasculature [4, 5, 12]. Strain magnitudes of 5-10% are considered physiological, whereas strains of 15-20% are characteristic of pathological situations [13–16]. Importantly, these mechanical forces are highly dynamic due to blood flow pulsatility. Although pulsatility has traditionally been assumed to be completely dampened by the time blood reaches the microvasculature, recent data challenge this consensus and have reported significant velocity and diameter oscillations even in the smallest capillaries [17–21]. Interestingly, in diseases such as hypertension, the higher pulse pressure penetrates deeper into the vascular tree, further increasing pulsatility in the microvasculature [7, 22].

Subjecting ECs to controlled levels of shear stress in vitro is readily accomplished in cone-and-plate systems, parallel plate flow chambers, and microfluidic channels [23]. However, these devices fail to stretch the cells. The effect of stretch on cells has traditionally been studied using planar uniaxial or biaxial stretching devices [24–26], but these systems do not incorporate flow and fail to capture the impact of substrate curvature, a particularly important consideration in the microvasculature. Microfluidic chips with pneumatic actuation in either planar [27–31] or curved [32,33] configurations have been reported, but these systems require external actuation and do not mimic the native substrate in terms of curvature, fibrillar structure, or stiffness. One platform that has been used to study the combined effects of shear stress and stretch on ECs is the inflatable vessel, where PDMS tubes are pressurized to generate both flow and circumferential strain [34–38]; however, the tubes have diameters of 5 mm, too large for studies of the microvasculature and incompatible with high resolution cellular imaging.

More recently, there has been mounting interest in vessel-on-chip systems that enable the application of controlled shear stress while providing a biologically relevant substrate and a realistic geometry [39–43]. A popular design consists of an endothelialized perfusable channel inside a collagen or fibrin hydrogel [44–49]. However, current microvessel-on-chip systems do not allow stretch due to the difficulty of incorporating actuators into micron-scale channels inside soft hydrogels. The ability to actuate hydrogels in microvessels-on-chip in a simple and controlled manner would enable the application of strain and would thus allow exploration of the combined effects of flow and stretch in these systems. Such a capability would significantly expand the experimental toolbox in the field of endothelial mechanobiology.

Here, we present a novel collagen hydrogel-based flow-actuated microvessel-on-chip that produces shear and strain simultaneously and overcomes the limitations of previous systems. We fully characterize the mechanical performance of the system and demonstrate the inherent coupling between shear and strain that can be accurately described within the poroelasticity framework. We show the ability to subject a confluent and functional EC monolayer to physiological and pathological levels of shear stress and strain under both steady and pulsatile conditions. Finally, we illustrate how different aspects of the design of the system can be tuned to modulate the shear-strain coupling, thus providing a highly versatile platform for elucidating the effects of a physiologically relevant mechanical stress environment on ECs.

## Materials and Methods

### Microvessel-on-chip fabrication

As depicted in Figure 1 and Supplementary Figure S1, the microvessel-on-chip system consists of a chamber that houses a 120 µm-diameter endothelium-lined channel embedded in a soft collagen hydrogel. The different components of the system are next described.

**Figure 1.**
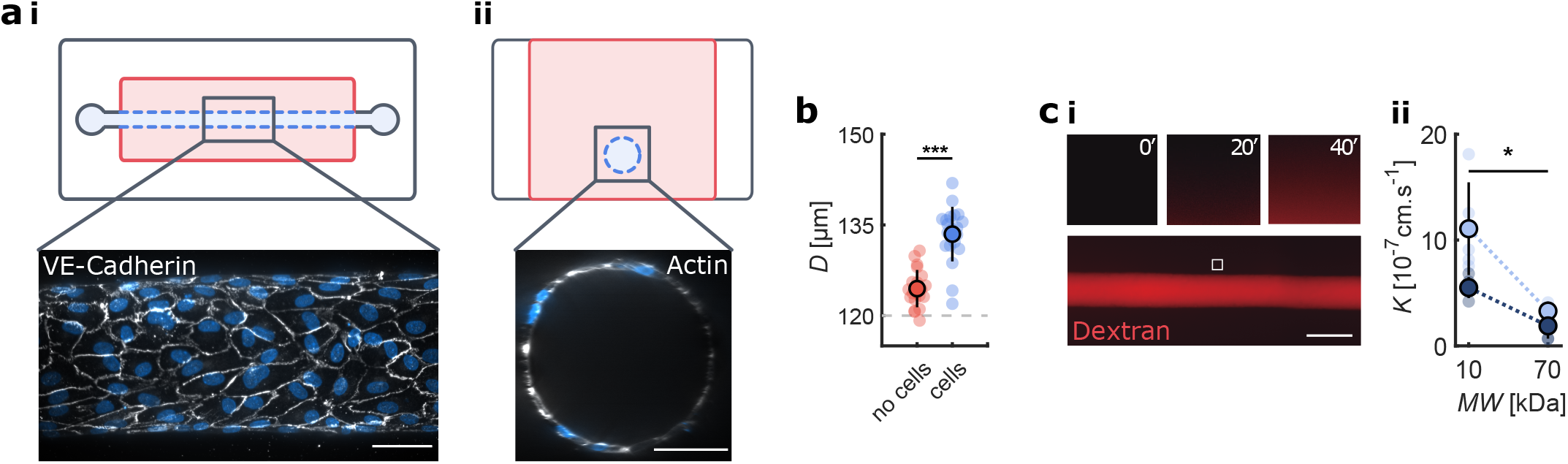
Endothelialization of the microvessel-on-chip. (**a**) Illustration of the microfluidic device (top view **i** and side view **ii**), showing the hydrogel (red), the channel (blue), and the PDMS housing (white). (**i**) Immunostaining of adherens junctions (VE-Cadherin, white) and cell nuclei (DAPI, blue), demonstrating the confluent monolayer lining the microvessel lumen. Scale bar 50 µm. (**ii**) Cross-section of the channel with actin staining in white (phalloidin) and cell nuclei in blue (DAPI), demonstrating the circularity of the endothelialized lumen. Scale bar 50 µm. (**b**) Microvessel diameter for bare gels (red) and with cells (blue). (**c**) **i** Diffusion of fluorescent dextran across the monolayer to quantify permeability. The three small panels show the progressive increase in dextran fluorescence intensity in the hydrogel at 0, 20, and 40 min. **ii** Quantification of monolayer hydraulic conductivity for two cell lines (Lonza, light blue; ScienCell, dark blue) and for two dextran molecular weights (MW), demonstrating barrier selectivity. Scale bar 150 µm. Dotted lines are guides for the eyes.

#### Housing chamber fabrication and preparation

The rectangular (15×2×3 cm) housing chamber was fabricated by pouring liquid elastomer mixed with 10% curing agent (polydimethylsiloxane (PDMS); Sylgard 184; Dow Corning) on a custom mold micromachined in brass to specific dimensions (Figure S2). After curing for 15 min at 180°C, the PDMS chamber was unmolded, and inlet and outlet ports were punctured using a hole punch. A 120 µm diameter acupuncture needle (Seirin) was introduced into the chamber, and liquid PDMS was used to fix it in place. With the needle in place, the bottom surface of the chamber frame was then bound to a coverslip through plasma activation. Finally, two fluid reservoirs, cut and punched from blocks of PDMS, were sealed with liquid PDMS to the inlet and outlet ports of the chamber, and the entire chip was then cured again for 1 h at 180°C. A critical feature is that the top of the chamber was maintained open.

The PDMS housing chamber was sterilized in 70% ethanol, dried, and plasma-activated for 45 s to render its surface hydrophilic and improve subsequent coating steps. The chamber was then covered with sterile PBS and placed under vacuum to remove bubbles from the inlet and outlet ports followed by 20 min of UV light exposure for final sterilization. To improve collagen adhesion to the PDMS walls, the chamber was coated with 1% polyethylenimine (PEI,an attachment promoter; Sigma-Aldrich) for 10 min followed by 0.1 % glutaraldehyde (GTA, a collagen crosslinker; Polysciences, Inc.) for 20 min.

#### Collagen hydrogel and microchannel fabrication

Collagen I was isolated from rat tail tendon as described previously [50]. Type I collagen solution was then prepared by diluting the acid collagen solution in a neutralizing buffer at a 1-to-1 ratio, pipetted into the housing chamber, and allowed to polymerize in a tissue culture incubator for 15 min for the baseline 6 mg/ml collagen concentration and for up to 4 h for lower collagen concentrations. The acupuncture needle was then carefully removed, and the needle holes were sealed with vacuum grease (Bluestar Silicones) to avoid leakage.

The hydrogel block formed inside the housing chamber was 15×2×3 mm (LxWxH); therefore, the length-to-diameter ratio of the 120 µm-diameter microchannel inside the chamber was 100. The microchannel was positioned far from all walls of the chamber: ∼400 µm from the bottom coverslip, 1 mm from the side walls, and more than 2 mm from the open top of the hydrogel (Figure 1.a, Figure S1).

#### Cell seeding and culture

Human umbilical vein ECs (HUVECs; Lonza) were cultured using standard protocols in Endothelial Growth Medium (EGM2; Lonza) and used up to passage 7. Upon confluence, HUVECs were detached from the flask using trypsin (Gibco, Life Technologies) and concentrated to 10^7^ cells.ml^-1^. 1 *µ*L of the concentrated cell suspension was pipetted through the inlet port of the device. The level of both reservoirs was adjusted to allow a very small flow from inlet to outlet in order to infuse the cell suspension into the channel. After a 5 min incubation, non-adhering cells in the channel were gently flushed out. The addition of 500 µL of medium to the inlet reservoir generated a small flow, necessary for cell survival and spreading. After 1 h, a flow rate of 2 µL min^*−*1^ was applied via a syringe pump (PhD Ultra, Harvard apparatus). A confluent monolayer was obtained in 36 h.

### Microvessel perfusion

Two different techniques were used to perfuse the microvessel at controlled flow rates: 1) a hydrostatic pressure head for the low flow (2 µL min^*−*1^) used for medium replenishment and waste product removal during long-term cell culture, and 2) syringe pump-driven flow at 10 or 50 µL min^*−*1^ used for the mechanical characterization of the microvessel and for the application of mechanical stimuli on the confluent HUVEC monolayer. To generate the hydrostatic pressure head, a PDMS reservoir was created by punching a 5 mm-diameter hole in a PDMS cubic block (1 cm on a side) and gluing the block to the inlet. The reservoir was continuously replenished using a syringe pump (Nemesys, Centoni). Using the reservoir as an intermediary between the syringe pump and the microchannel was particularly useful for preventing air bubbles within the channel. For the syringe pump-driven flow, the pump was connected directly to the microchannel inlet.

### Immunostaining

Cell-cell junctions were stained using a rabbit anti-VE-cadherin primary antibody (Abcam). Actin filaments and nuclei were stained using Alexa Fluor phalloidin (Invitrogen, Thermo Fisher Scientific) and DAPI (Sigma-Aldrich), respectively. Immunostaining was performed by slow infusion of reagents into the microchannel. Cells were fixed in 4% paraformaldehyde (PFA; Thermo Fisher Scientific) for 15 min, rinsed with phosphate-bufered saline (PBS), and then permeabilized with 0.1% Triton in PBS for another 15 min. The channel was then perfused with a 3% bovine serum albumin (BSA) solution in PBS for 1 h to block non-specific binding. Cells were incubated with the VE-cadherin primary antibody (1:400) in PBS for 1 h at room temperature and then rinsed with PBS for an additional 1 h. The channel was then perfused with an anti-rabbit secondary antibody (1:400), phalloidin (1:200), and DAPI (1:1,000,000) solutions. Finally, the cells were incubated overnight in PBS at 4°C. Samples were imaged using the NIS-Elements software on an epifluorescence inverted microscope (Nikon Eclipse Ti) and/or a Crest X-Light confocal system mounted on an inverted microscope (Nikon Eclipse Ti).

### Permeability measurement

After monolayer formation, HUVECs were maintained in culture for an additional 24 h under a 2 µL min^*−*1^ flow to allow cell-cell junction maturation. Rhodamine-dextran (10 and 70 kDa molecular weight; Life technologies) was used as a fluorescent tracer to assess monolayer macromolecular permeability. The cells were maintained at 37°C during permeability measurements by using a microscope equipped with a stage-top incubator.

50 µL of medium containing 2 µL of dextran was added to the inlet reservoir. The fluid volume in the outlet reservoir was adjusted to 50 µL to avoid the creation of a hydrostatic pressure difference. The open top of the chip was covered with PBS during image acquisition to avoid evaporation in the hydrogel.

Images of dextran diffusion into the hydrogel were acquired at 4 frames.min^-1^ for 2 h and at several positions along the length of the channel using a Flash 4.0 CCD camera (Hamamatsu) mounted on an inverted fluorescence microscope (Nikon eclipse Ti) equipped with a 10x objective. The intensity profiles from the images were extracted using ImageJ. The evolution of dextran intensity over time was measured in the channel lumen and in the gel at a radial position 15 µm from the microvessel walls.

As in previous work [51], the hydraulic conductivity (which is directly proportional to permeability) (*K*) of endothelialized channels to dextran was calculated as follows:

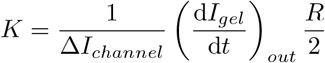

Where 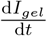 is the rate of change of fluorescence intensity in the hydrogel, Δ*I*_*channel*_ is the increase in fluorescence intensity within the lumen (due to luminal filling with dextran), and *R* is the microvessel radius.

For bare (non-endothelialized) channels, dextran (both 10 and 70 kDa) diffuses virtually instantaneously into the gel after entering the channel. Therefore, hydraulic conductivity values in that case cannot be computed using this same approach.

### Particle tracking velocimetry

Particle tracking velocimetry (PTV) was used to determine the velocity profile within the microvessel.

The channel was perfused with a suspension of (1.7 µm-diameter microbeads (Polysciences, Inc.) in PBS (1.8×10^9^ beads.mL^-1^) at controlled flow rates ranging from 10 to 50 µL min^*−*1^. Images were acquired with a high speed camera (Proton Fastcam SA3) and a 40x objective at 5,000 to 20,000 frames per second (fps). The cameraïs field of view was sufficiently large to include the entire channel diameter and 50 µm of channel length. The acquisition rate was enough for more than 500 beads (on average) to pass through, allowing precise velocity profile reconstruction. The images were pre-processed to remove the stationary background (by subtracting the average image) and inverted to make beads white. The tracking was performed using TrackMate, a free ImageJ plugin [52]. The beads (7 pixels in diameter) were detected automatically and linked in a continuous track by a linear motion tracker. Finally, erroneous tracks were removed with two low pass filters: velocity standard deviation and mean velocity.

For steady flow, the mean radial position and mean axial velocity were computed for each particle track. Because the illumination plane had a certain thickness (estimated to be 40 µm), the velocities at each radial position were dispersed. Because the particles of interest are the ones in the channel mid-plane, only beads having velocities within 5-10% of the maximum value at each radial position were used for velocity profile reconstruction. The experimental points were fitted to a parabola, in accordance with the Poiseuille flow profile for fully developed steady flow in a cylindrical channel at low Reynolds number (Re ∼ 10).

For pulsatile flow, the processing was adapted to access the instantaneous flow profile (Movie 6). The recorded movie was cut into hundreds of short time segments to ensure a nearly constant velocity during each segment. The mean axial velocity was computed for individual tracks in each segment. The highest velocities representing the particles passing close to the channel centerline were selected to obtain the maximum velocity as a function of time. At this stage, the reconstructed profile was not sufficiently precise to yield the flow rate directly since very few particles were present in each time segment (and usually none close to the channel walls), leading to a large uncertainty in the measured channel radius. The instantaneous radius was therefore extracted from the live recording of channel wall movement during the pulsatile cycle. Assuming a time-dependent parabolic flow profile, expected to be accurate for the low Womersley number present here

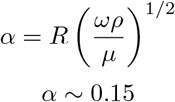

and verified to be the case by plotting instantaneous velocity profiles (Movie 6), the mean flow rate and wall shear stress were computed for each time segment based on the maximum velocity (from PTV) and the radius (from channel wall tracking) using Poiseuille’s equation for flow in a circular cross-section channel.

### Stretch measurement

An increase in the luminal pressure leads to a pressure difference across the wall and subsequent channel dilation and circumferential strain. The circumferential strain was defined as the ratio of the increase in perimeter of the cross section of the channel to the initial perimeter. Because the perimeter depends linearly on diameter, the circumferential strain (when homogeneous) is equal to the normalized diameter increase. The circumferential strain was therefore obtained directly from the diameter change. Two methods were used to compute the strain: manual measurement of the diameter change for the case of constant strain and automated diameter tracking for cyclic strain. The manual measurements of channel diameter were performed in ImageJ, first on an image of the channel before the pressure increase and then on an image taken a few seconds after the pressure increase. The automated tracking of channel diameter was conducted using Clickpoints, an open-source tracking software [53].

### Optical coherence tomography imaging

To visualize the channel cross section, Optical coherence tomography (OCT) images were acquired using a Ganymede OCT system (Thorlabs). The acquisition speed was 37 fps.

### Statistical analysis

For Figures 1-4, all data are plotted as mean ± SD. An unpaired Student t-test was used for significance testing between two conditions. Statistical tests were performed using Matlab. For each condition, at least three independent microvessel systems (5 regions per system) were used for the imaging and data analysis. **** denotes p *<* 0.0001, *** denotes p *<* 0.001, ** denotes p *<* 0.01, and * denotes p *<* 0.05.

## Results

### Cellular coverage and monolayer integrity

Human umbilical vein ECs (HUVECs) seeded in the microvessel-on-chip and subjected to a low flow of 2 µL min^*−*1^ for 48 h develop into a confluent monolayer with clearly delineated adherens junctions as demarcated by VE-cadherin staining (Figure 1a). Despite the softness of the collagen hydrogel within which it is embedded, the microvessel lumen has a circular cross-section as determined by 3D reconstruction of confocal microscopy images (Figure 1aii, Movie 1) and OCT imaging (Figure S1). Consistent with the size of microneedle used in its fabrication (nominal diameter of 120 µm), the microchannel prior to cell seeding has a diameter of 125±5 µm. Interestingly, this diameter increases to 134±5 µm after endothelial monolayer formation (Figure 1b).

Fluorescently labeled dextran infused into the microvessel lumen results in dextran transport across the endothelial monolayer and into the collagen hydrogel over a period of minutes (Figure 1c). The hydraulic conductivity (proportional to the permeability) of the microvascular wall is computed (see Methods) for dextran of two different molecular weights (10 and 70 kDa) and using HUVECs from two commercial sources (Lonza and Sciencell). For cells from both sources, the hydraulic conductivity to 70 kDa dextran is significantly lower than that for 10 kDa dextran (Figure 1c), demonstrating the ability of the endothelium to discriminate among molecules of different sizes and thus to act as a selective permeability barrier. At each molecular weight, there is a 2 to 3-fold difference in computed hydraulic conductivity between the cells from the different sources, underscoring the importance of carefully characterizing the cells and the potential perils of mixing cells from different commercial sources. The hydraulic conductivity values obtained here are consistent with measurements published in other in vitro microvessels [54–56].

### Wall shear stress in the microvessel: interplay between luminal flow and gel porosity

Particle tracking velocimetry (PTV) measurements (Figure S3, Movie 2) confirm that for steady flow, the velocity pro-file inside the microvessel is parabolic, consistent with the Poiseuille flow expected for the range of Reynolds numbers studied (2 to 10) (Figure 2a,b). Adjusting the inlet flow rate allows fine tuning of the wall shear stress magnitude from 0.1 to 2 Pa (Figure 2c), spanning the physiological range in the microvasculature.

**Figure 2.**
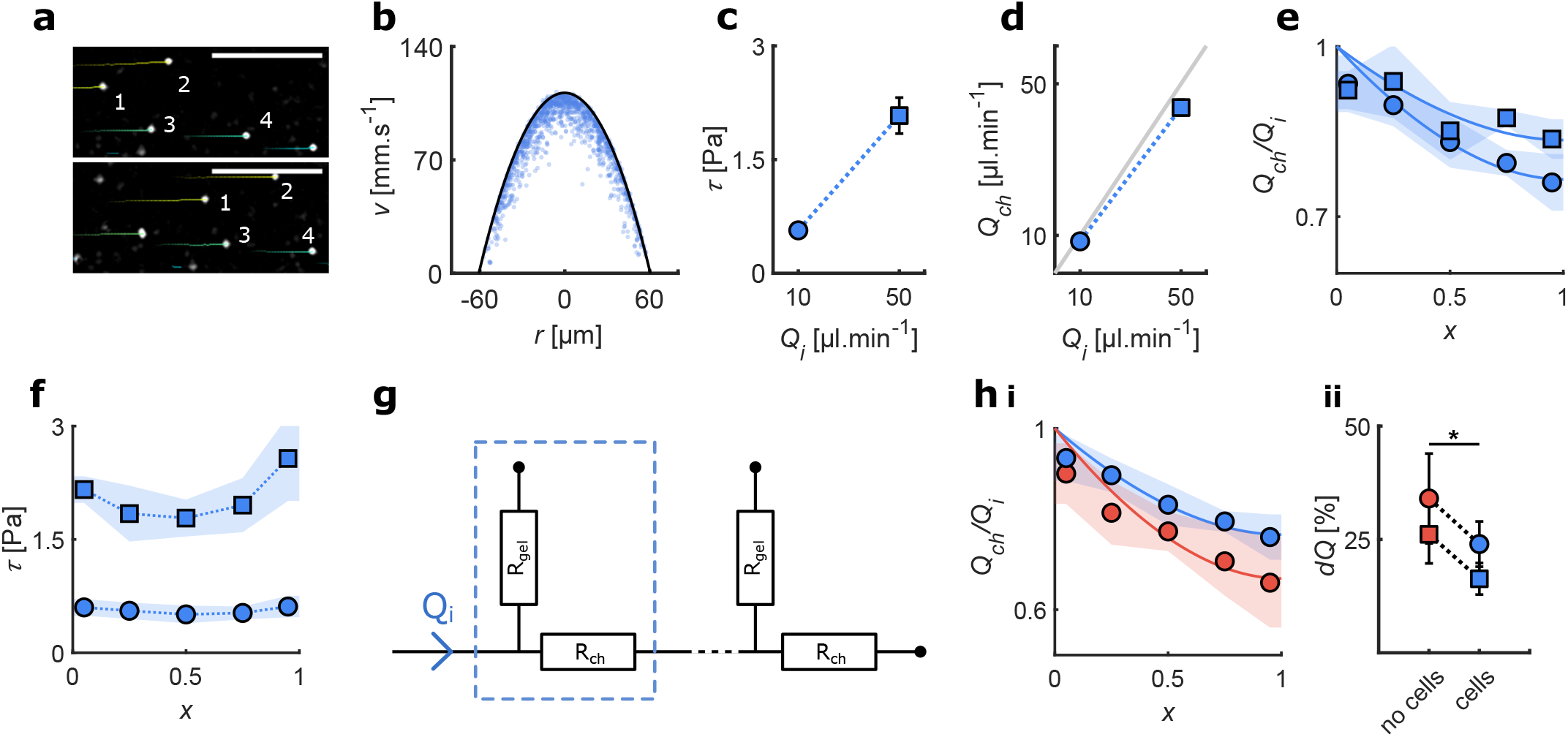
The imposed flow rate controls the wall shear stress. Blue represent data from endothelialized channels and red from bare channels. (**a**) Particle tracking velocimetry at two successive time points. (**b**) Velocity profile in the channel midplane; each dot represents the average velocity of one microbead. A parabola is fitted to the 5% fastest beads to measure the flow rate and wall shear stress. (**c**) Measured wall shear stress increases with flow rate. (**d**) The measured flow rate in the channel (symbols) is smaller than the imposed flow rate (solid gray line) due to fluid seepage into the gel. (**e**) Normalized flow rate in the channel at different axial positions for the two different imposed flow rates. The solid line is the best fit of the analytical model. (**f**) Wall shear stress as a function of position for the two different imposed flow rates. (**g**) Schematic of the equivalent electrical circuit used for the analytical modeling. *R*_*gel*_ is the hydraulic resistance of the porous gel and *R*_*ch*_ is the hydraulic resistance of the cylindrical channel. (**h**) **i** Flow loss is larger in bare channels (red) compared to channels with an endothelial monolayer (blue); *Q*_*i*_ = 10 µL min^*-*1^. Solid lines indicate the predictions of the analytical model. **ii** Flow loss in a bare gel G (red) and with cells C (blue), measured at the two different imposed flow rates. In all panels circles correspond to *Q*_*i*_ = 10 µL min^*-*1^ and squares correspond to *Q*_*i*_ = 50 µL min^*-*1^. Dotted lines are guides for the eyes.

The measured flow rate in the microvessel averaged over the entire length *Q*_*ch*_ is lower than the imposed flow rate at the inlet *Q*_*i*_: 8.4 vs. 10 µL min^*−*1^ and 44 vs. 50 µL min^*−*1^ (Figure 2d). This difference is attributable to a progressive decrease in luminal flow rate with axial position (Figure 2e). This luminal flow loss is due to the hydrogel porosity whereby part of the flow seeps into the gel surrounding the channel towards the open top which acts as a low pressure outlet. Interestingly, the wall shear stress does not exhibit a similar decrease (Figure 2f) due to the axial variations in channel diameter (see next section) that compensate the flow loss.

#### Understanding the division between luminal and transmural flow based on electrical circuit analogy

A useful analogy to understand the competition between the two flow paths, flow in the lumen and flow through the gel, is the equivalent electrical circuit (Figure 2g). In this analogy, the electrical current and the potential difference driving this current are analogous to the flow rate and pressure gradient, respectively. Because the channel here is much longer than wide, it can be modeled by an infinite number of elementary units along the axial position. At any axial position *x*, the flow splits between two paths: a microchannel path with resistance R_ch_ that depends on the channel radius and a gel path with resistance R_gel_ that depends on hydrogel dimensions and concentration. Thus, the remaining flow rate in the channel at *x* +*dx* is smaller than that at *x*. From this equivalent electrical circuit, an analytical model of the free and porous medium flow through the microvessel is derived (see Appendix A) and is shown to accurately predict the flow drop in the channel for a bare (no cells) gel (Figure 2hi, red line).

#### Cell monolayer acts as a flow barrier

To investigate the role the endothelium might play in altering the microchannel flow loss, *Q*_*ch*_ is also measured in bare channels containing no ECs. The flow loss is significantly larger in bare gels than in endothelialized channels (Figure 2h), indicating the confluent monolayer acts as a semi-permeable membrane that reduces fluid leakage into the porous gel. The equivalent circuit model can also account for the contribution of the endothelium by modeling the monolayer as an additional resistance between the channel and gel. To this end, the gel permeability is first obtained by fitting the analytical flow drop to experimental data from bare channels (Figure 2hi, red line). The additional hydraulic resistance due to the cell monolayer is then obtained by fitting the analytical flow drop to experimental data from endothelialized channels using the gel permeability found above. The model correctly captures the decrease in flow loss due to the presence of the cell monolayer (Figure 2hi, blue line). These results confirm the validity of a simple equivalent circuit analysis and underscore the importance of taking porous medium flow into account to understand the mechanical environment inside hydrogel-based microvessel-on-chip systems.

### Circumferential strain and gel elasticity

A principal novelty of the system presented here is its ability to produce controlled and physiologically relevant circumferential strains. This is accomplished through luminal flow actuation whereby increasing the flow rate in the microchannel increases luminal pressure, which compresses the soft hydrogel, dilates the channel, and stretches the endothelial monolayer (Figure 3a, Movie 3). OCT imaging demonstrates that the channel remains nearly circular during dilation, leading the cells to experience a circumferentially uniform strain field (Figure 3aii). When the flow is arrested, the luminal pressure rapidly drops, and the channel relaxes back to its original diameter. The hydrogel deformation is perfectly reversible, characteristic of elastic solids (Figure 3a), an essential feature to ensure controllable and robust actuation. The circumferential strain *ε*, quantified as the ratio of change in diameter to the initial diameter, reflects this reversible behavior (Figure 3aiii). The changes in luminal flow rate that drive wall strain also lead to changes in wall shear stress (Figure 3aiii), demonstrating the coupling between shear and strain in our system. As the strain ensues from luminal pressure, changing the imposed flow rate at the microvessel inlet *Q*_*i*_ modulates the level of applied strain. Average strains of 4% and 12%, spanning the relevant physiological range, are obtained as *Q*_*i*_ is increased from 10 to 50 µL min^*−*1^ (Figure 3bi).

**Figure 3.**
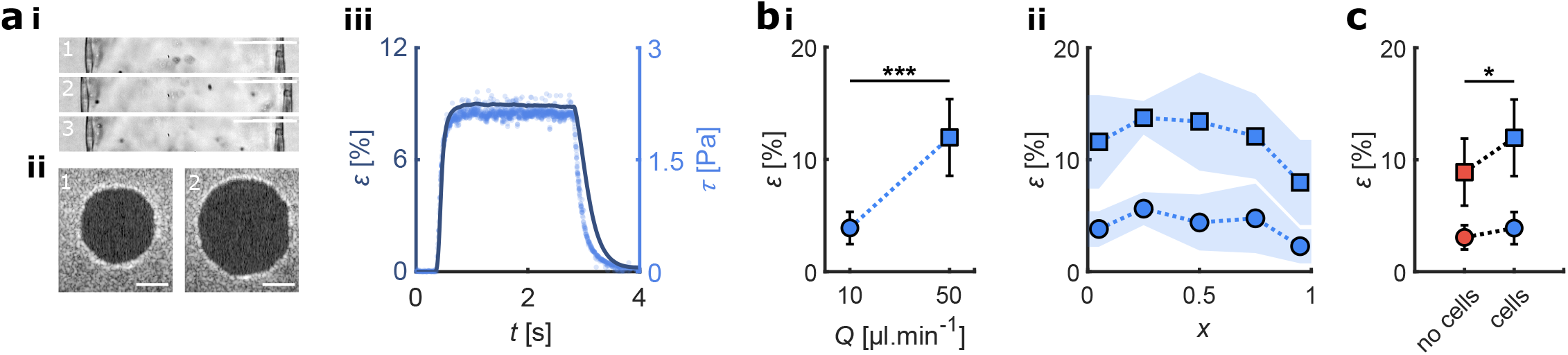
The imposed flow rate controls the circumferential strain. (**a**) Channel dilation following a step in flow rate. **i** Brightfield image of the channel’s midplane before (1), during (2) and after (3) the step. **ii** OCT image of the channel cross-section before (1) and during (2) the step. **ii** Dynamics of channel dilation: strain *ε* (dark blue, solid line) and wall shear stress *τ* (light blue, dots) as a function of time during a flow step. Scale bars 50 µm. (**b**) Strain as a function of the imposed flow rate. **i** Average strain for two imposed flow rates. **ii** Strain as a function of position along the length of the channel for two imposed flow rates. (**c**) Strain for a bare gel (red) and a channel lined with an EC monolayer (blue) for two imposed flow rates. In panels b and c, circles correspond to *Q*_*i*_ = 10 µL min^*-*1^ and squares correspond to *Q*_*i*_ = 50 µL min^*-*1^. Dotted lines are guides for the eyes.

For a given *Q*_*i*_, the microvessel strain *ε* depends on the axial position. The deformation is small close to the two ends of the microvessel because the gel is chemically bound to the PDMS housing, imposing a zero displacement at the wall (Figure 3bii). Away from the microvessel ends, *ε* decreases axially as a result of the hydraulic pressure drop created by the flow. To investigate the possible impact of the endothelial monolayer on microvessel strain, the average strains are also measured in bare (no cells) channels at flow rates of 10 and 50 µL min^*−*1^. Strain levels are significantly higher in the presence of an endothelium (Figure 3c). Although somewhat counter-intuitive, the increased deformation is explained by the flow barrier effect of the cell monolayer and the poroelastic properties of the hydrogel. More specifically, the presence of the EC monolayer decreases fluid seepage into the gel which leads to a lower gel pressure (while the pressure in the channel remains virtually unchanged). In poroelastic materials, pore deformation is coupled to pore pressure, with increased pore pressure leading to reduced material compression. Thus, the lower gel pressure in the presence of an endothelium allows increased gel deformation and higher strains. This phenomenon illustrates the coupling between the flow rate and strain due to gel poroelasticity.

### Tuning the shear-strain coupling

As mentioned in previous sections, the luminal flow rate *Q*_*ch*_ and the channel strain *ε* are coupled through pressure. Additionally, the wall shear stress *τ* depends not only on *Q*_*ch*_ but also on channel radius, which is modified by the strain *ε*. As a result, the shear stress *τ* and the strain *ε* are tightly coupled and vary as functions of the imposed flow rate *Q*_*i*_, the hydraulic resistances, and the spring constant of the gel. To tune the coupling between shear stress and strain, three different strategies are explored.

#### Strategy 1: Modifying outlet resistance to control channel pressure

The simplest method to increase the channel pressure is to add hydraulic resistance to the outlet, *R*_*out*_. Tubing of creased from 0 to 12×10^12^ N.s.m^-5^ (corresponding to outflow tubing length increasing length connected to the outlet is used to progressively increase the outflow resistance *R*_*out*_ (Figure 4a.i). Flow loss, strain and shear stress are then measured for each *R*_*out*_ in the same chip for *Q*_*i*_ = 10 µL min^*−*1^. As *R*_*out*_ is increasing from 0 to 9 cm), luminal pressure also increases, leading to an increase in average flow loss into the hydrogel (Figure 4a.ii) as well as an increase in average channel strain (Figure 4a.iii). These two effects lead to a reduction in average wall shear stress (Figure 4a.iv). These results demonstrate a strong coupling between channel strain and wall shear stress (Figure 4a.v). As for the axial variations (Figure S4a), the larger flow loss into the gel at high *R*_*out*_ translates into significantly larger axial gradients of wall shear stress. Interestingly, the strain slope remains fairly constant with *R*_*out*_ and is simply offset for the different outflow resistance values.

**Figure 4.**
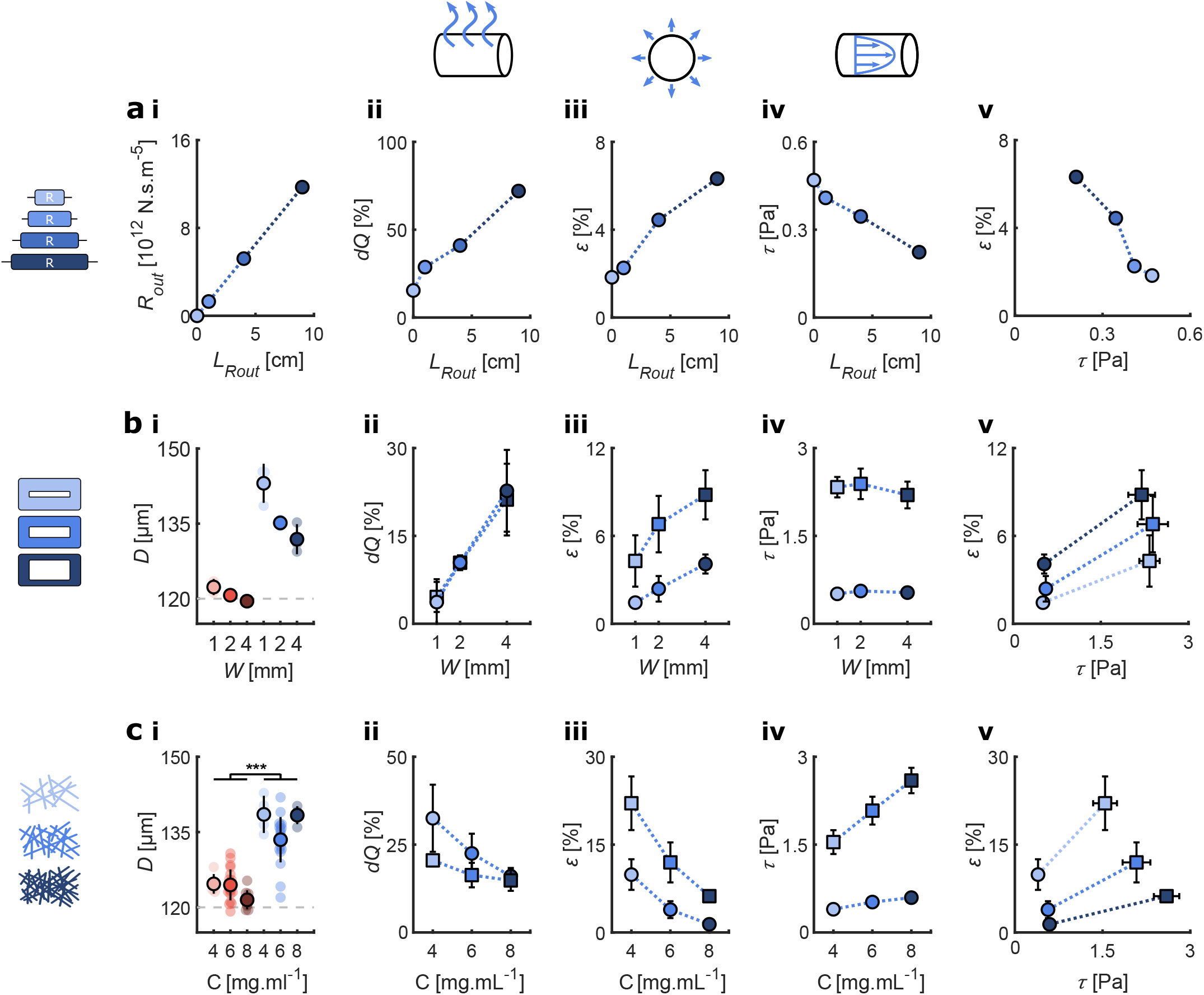
Three strategies to modulate the shear-strain coupling: outlet resistance, gel width and gel concentration. (**a**) Increasing the outlet resistance by increasing the length of the outlet tubing *L*_*Rout*_. All results are for *Q*_*i*_ = 10 µL min^*-*1^. **i** Average outlet resistance. **ii** Average flow loss in the channel *dQ*. **iii** Average wall strain *ε*. **iv** Average wall shear stress *τ*. **v** Strain-wall shear stress coupling. (**b**) Changing the gel width. All results are repeated for two imposed flow rates. **i** Average channel diameter *D*. **ii** Average flow loss in the channel *dQ*. **iii** Average wall strain *ε*. **iv** Average wall shear stress *τ*. **v** Strain-wall shear stress coupling. (**c**) Changing the gel concentration. All results are repeated for two imposed flow rates. **i** Average channel diameter *D*. **ii** Average flow loss in the channel *dQ*. **iii** Average wall strain *ε*. **iv** Average wall shear stress *τ*. **v** Strain-wall shear stress coupling. In all panels, circles correspond to *Q*_*i*_ = 10 µL min^*-*1^ and squares to *Q*_*i*_ = 50 µL min^*-*1^. Red indicates bare gels and blue endothelialized channels. Dotted lines are guides for the eyes.

#### Strategy 2: Modifying gel width to change its hydraulic resistance and spring constant

Another strategy to tune the shear stress-strain coupling is to change the width of the chamber housing the hydrogel, which modulates both the hydraulic resistance and spring constant of the gel. Flow loss, strain, and wall shear stress in the channel are measured for chamber widths of 1, 2, and 4 mm at *Q*_*i*_ values of 10 and 50 µL min^*−*1^ (Figure 4b). Although hydrogel width does not change the diameter of bare channels, endothelialized channels in narrower hydrogels have larger diameters (Figure 4b.i). Because wider hydrogels have smaller *R*_*gel*_, increasing gel width leads to significantly increased flow loss (Figure 4b.ii). The smaller spring constants in wider hydrogels lead to higher gel deformation and thus to larger strains (Figure 4b.iii). Interestingly, the average wall shear stress remains relatively constant because of the smaller initial diameter found in wider hydrogels that compensate both the increased flow loss and increased strain (Figure 4b.iv). Therefore, this strategy provides a pathway for largely decoupling wall strain from wall shear stress (Figure 4b.v). As for the axial variations (Figure S4b), the strain slope is again maintained fairly constant and is simply offset for the different gel widths, while the wall shear stress does not exhibit significant axial variations.

#### Strategy 3: Modifying gel concentration to change its permeability and stiffness

The third strategy to tune the coupling between shear stress and strain is to modify hydrogel concentration which determines both gel permeability and stiffness. The flow rate, strain and wall shear stress in the channel are measured for the three collagen densities of 4, 6 and 8 mg mL^*−*1^ and at the two *Q*_*i*_ values of 10 and 50 µL min^*−*1^. Hydrogel concentration does not change the baseline diameter of either bare or endothelialized channels (Figure 4c.i). Because denser gels are less permeable and stiffer, channels exhibit smaller flow loss and smaller average strain (Figure 4c.ii & iii). The two effects combine to significantly increase the average wall shear stress in denser gels (Figure 4c.iv). Thus, using this strategy preserves the coupling between wall strain and wall shear stress (Figure 4c.v), although this coupling is somewhat weaker than that associated with strategy 1. The shape of the axial variations of both the strain and the wall shear stress are unchanged by varying gel concentration and are simply offset as gel concentration changes (Figure S4c).

The quality and homogeneity of the collagen hydrogel impacts its mechanical characteristics. The difference between two collagen batches of the same concentration was investigated. Batch #1 was visibly less homogeneous than batch #2: dissolution required a longer time, fibers were observed under the microscope, and numerous holes and defects were visible with OCT imaging. Predictably, batch #1 led to significantly higher flow loss and average strain than batch #2 (Figure S5), a behavior similar to that of a lower gel concentration, underscoring the sensitivity of the actuation to the quality of the gel. Consequently, throughout the present study, all the chips used in one experiment were fabricated with the same batch of collagen in order to ensure that only the tested parameter is changed among replicates.

#### Comparing strategies: an operating manual for the microvessel-on-chip

Integrating the information gleaned from the three different strategies described above provides a form of “operating manual” for the microvessel-on-chip system studied here, as schematically illustrated in Figure 5 which depicts the relative change in strain and wall shear stress afforded by each strategy. From a practical standpoint, changing the outlet resistance is by far the simplest strategy since it is straight-forward, highly reproducible, and can be implemented in the same chip. Contrarily, changing hydrogel concentration and gel width require modifications to the fabrication and thus different chips.

**Figure 5.**
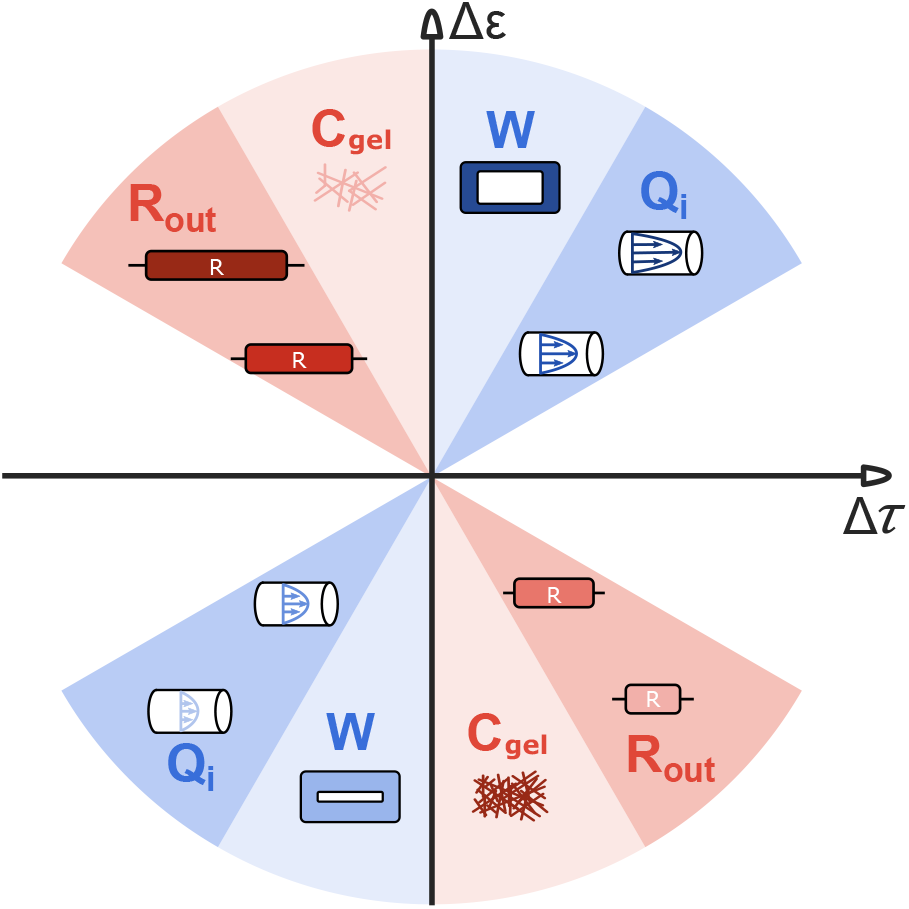
Schematic depiction of the relative effects of changing the outlet resistance (*R*_*out*_), gel concentration (*C*_*gel*_), gel width (*W*), and imposed flow rate *Q*_*i*_ on the coupling between wall strain and wall shear stress. The origin denotes the baseline conditions. Positive values of Δε and Δτ respectively denote increases in strain and wall shear stress relative to the baseline, while negative values correspond to decreases relative to the baseline. The colored wedges describe each of the tuning strategies with the blue and pink colors respectively indicative of strategies that have a similar or opposite effect on strain and wall shear stress. The position of each wedge describes the relative effect of the strategy it represents on strain and wall shear stress. For instance, changing gel width or gel concentration has a larger effect on channel strain than on wall shear stress, whereas changing outlet resistance or inlet flow rate has relatively similar effects on both strain and wall shear stress.

How can the information from the different tuning strategies be exploited as an operating manual and when might each of these strategies be most useful? Because modifying the outlet resistance has opposite effects on strain and wall shear stress (Figure 5), it constitutes an effective strategy for elucidating if a particular cellular response is driven more by strain or by shear stress. Furthermore, because manipulating outlet resistance dramatically impacts the amount of flow loss into the gel, it generates significant gradients in wall shear stress along the channel length. Therefore, this strategy would also be particularly useful for studies that aim to elucidate the effect of spatial gradients of shear stress on the endothelium. Modulating the gel width allows a wide range of strains while maintaining the wall shear stress relatively constant (Figure 5), thereby providing a roadmap for separating the effects of shear stress and strain. Finally, changing gel concentration allows a wide range of strains and shear stresses in a manner similar to manipulating out-flow resistance, with smaller effect on the wall shear stress (Figure 5). An additional consideration, however, is that gel concentration has a significant impact on gel stiffness; therefore, in comparison to controlling outflow resistance, manipulating gel concentration would also provide the capability of studying EC responses to changes in substrate stiffness. In addition to the effects of these three strategies, Figure 5 also illustrates the effect of simply changing the imposed flow rate. Increasing the inlet flow rate increases both wall strain and wall shear stress and thus provides the full coupling that might be most representative of the situation encountered in vivo.

### Model of the channel and gel flow

The analytical model derived from the equivalent electrical circuit analogy is detailed in Appendices A and B. This model, which considers free fluid flow within the channel and porous medium flow governed by Darcy’s law within the hydrogel, can be used to demonstrate how the shear-strain coupling is impacted by the three tuning strategies and to guide the decision making process of microvessel-on-chip users. As detailed in the following section, modifications of the experimental system are implemented in the model by varying the constitutive parameters.

To model the impact of the added outlet resistance *R*_*out*_, the boundary condition for the outlet pressure is modified to the following:

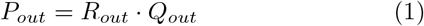

First, the gel permeability, the only unknown parameter of the system, is obtained by fitting the model to the experimental flow drop measured for *R*_*out*_ = 0. Then, the flow rate *Q*_*ch*_ and the pressure P at the different axial positions are calculated for different values of *R*_*out*_. Our model correctly predicts the larger flow loss *dQ* for increasing *R*_*out*_ and matches the experimental data closely (Figure 6a.i). It is important to note that *dQ* saturates at high *R*_*out*_. Indeed, when *R*_*out*_ is large, *Q*_*out*_ is small due to the large flow loss, leading in turn to a reduced effect of *R*_*out*_ on the luminal pressure. Additionally, the ratio of experimental strain to computed channel pressure is found to be constant (Figure 6a.ii). This proportionality constant is related to the Young’s modulus of the gel which, by definition, relates stress to strain.

**Figure 6.**
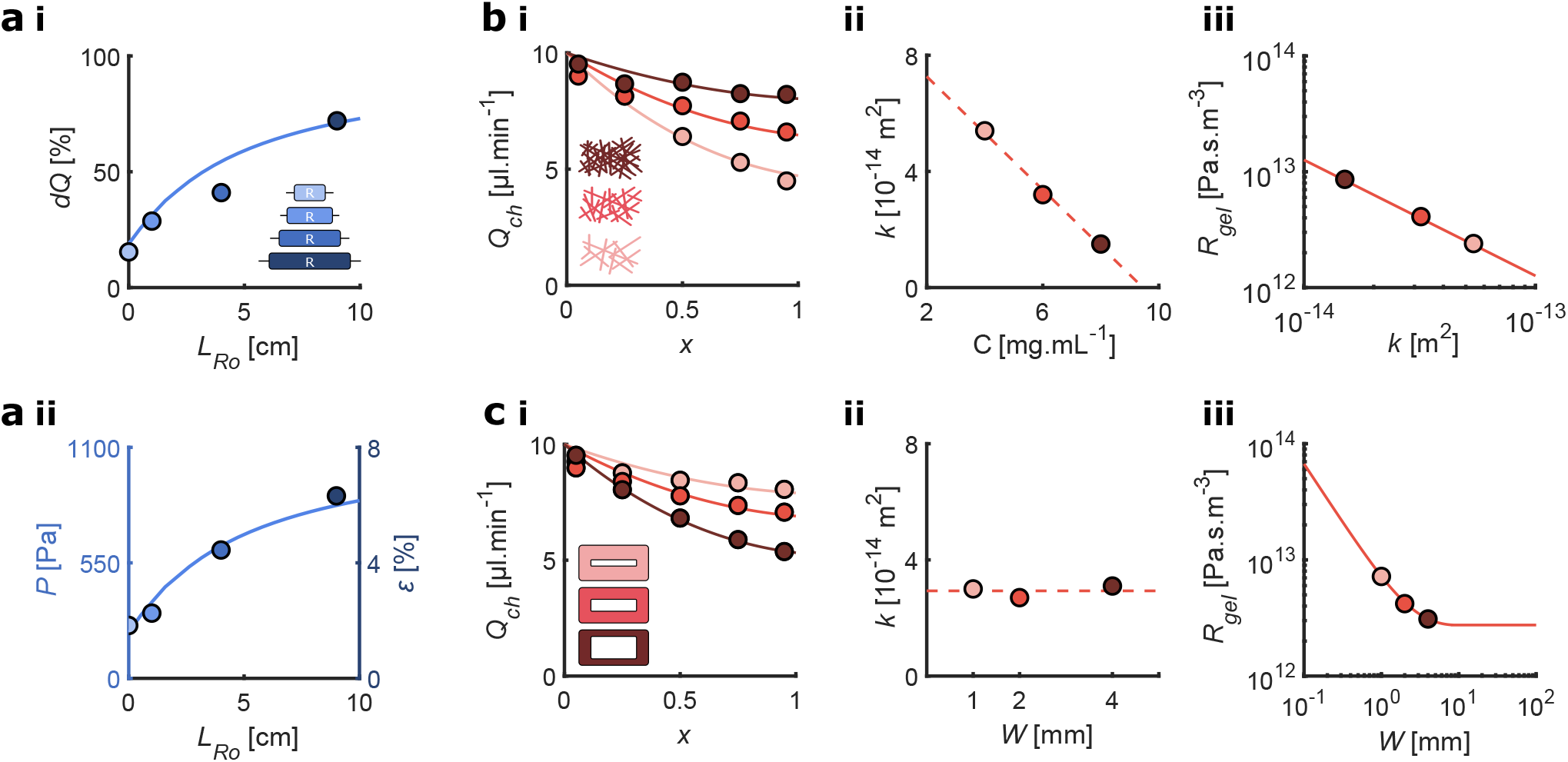
Analytical model of the porous medium flow based on the equivalent electrical circuit. (**a**) Modeling the impact of outlet resistance (darker colors correspond to increasing resistances, the cell monolayer is included in all cases). **i** Predicted flow loss and measured flow loss as a function of outlet resistance length. The model curve (solid line) is based on the permeability fitted to the baseline condition (*R*_*out*_=0). **ii** Predicted luminal pressure (solid line) and measured strain (dots) as a function of outlet resistance. The model curve (solid line) is based on the permeability fitted to the experiment with no outlet resistance. (**b**) Modeling the impact of hydrogel width (light, medium and dark red correspond to 1, 2, and 4 mm-wide bare gels, respectively). **i** Channel flow as a function of the axial position: visualization of the least squares fitted curve to the experimental data for the optimal hydrogel width. **ii** Dotted line shows hydrogel permeability found through the fit as a function of hydrogel width. Circles represent experimental data (darker colors correspond to increasing width). **iii** Hydraulic resistance of the hydrogel as a function of hydrogel width. (**c**) Modeling the impact of hydrogel concentration (light, medium and dark red correspond to 4, 6, and 8 mg.mL^-1^ bare gels, respectively). **i** Channel flow as a function of the axial position: visualization of the least squares fitted curve to the experimental data for the optimal hydrogel concentration. **ii** Hydrogel permeability, found through the fit, as a function of hydrogel concentration. **iii** Hydraulic resistance of the hydrogel as a function of hydrogel permeability. In all panels, circles correspond to experimental data or values inferred from experimental data. Solid lines are the predictions of the model and dashed lines are linear fits.

To elucidate the effect of variations in gel width and concentration, the layer of cells is not taken into account. Only the gel is modeled, and the experimental data from bare gels are used. Theoretical curves of *Q*_*ch*_ as a function of axial position are fitted to the experimental flow drop for the three gel densities (Figure 6b.i). The optimal gel permeability *k*_*gel*_ for each condition is found by least squares fitting. The fitted gel permeability values are 5.4, 3.2, and 1.6 × 10^−14^ m^2^ for gel densities of 4, 6, and 8 mg mL^−1^, respectively. The model thus correctly predicts a smaller permeability for the denser gels. The permeability of other gels can be estimated by extrapolating the empirical linear *k*_*gel*_ - *C*_*gel*_ relationship (dashed line), provided their concentration is of the same order of magnitude (Figure 6b.ii). The gel resistance *R*_*gel*_ is inversely proportional to the gel permeability, as predicted by Darcy’s law (Figure 6b.iii). As such, denser gels have larger resistances, leading to smaller flow losses.

The same approach is used to understand the role of gel width on porous medium flow in the system. Gel resistance *R*_*gel*_ depends not only on gel permeability but also on its dimensions. The geometry is defined by the cross-section of the system: a large rectangle with a small circular hole inside of it, in which fluid transport occurs from the hole towards the open top (zero pressure). Because the transport does not take place only in one dimension, there is no analytical expression for the gel resistance. The gel is therefore modeled as two resistances in series, assuming first radial flow across a torus, then longitudinal flow across a rectangle (see Appendix). Defined as such, the total gel resistance can be expressed analytically using Darcy’s law as a function of the three relevant length scales: the radius of the channel and the width and height of the gel. The analytical solution is fitted to experimental data for the three different gel widths to obtain the permeability values, all found to be around 3×10^−14^ m^2^ (Figure 6c.i,ii). Indeed, as the gel concentration is identical for all widths, the permeability is expected to remain unchanged. The model thus correctly predicts the different flow drops only through the change in gel width, included in the expression for *R*_*gel*_ (Figure 6c.i,iii). Plotting the analytical *R*_*gel*_ as a function of gel width W shows two regimes (Figure 6c.iii). The three experimental designs (dots) are in the transition zone. When the gel is narrow, most of the resistance comes from the longitudinal transport: *R*_*gel*_ is inversely proportional to the width W. When the gel is wide, the resistance becomes constant: all the flow crosses the torus whose thickness is the constant height of the gel. In the transition zone, the contribution of the torus is significant, and *R*_*gel*_ varies logarithmically with gel width.

The analytical model of the free and porous medium flow, based on the equivalent electrical circuit, correctly replicates the experimental observations for three sets of variations: boundary conditions (outlet resistance), geometrical parameters (gel width), and constitutive parameters (gel concentration). Flow drop variations are reproduced and interpreted, and the computed luminal pressure, which cannot be measured experimentally, explains the observed strains. Although not presented here, wall shear stress can also be computed from the analytical flow values by substituting the deformed geometries based on the experimental measurements of strain into the model. The model can predict the behavior of new experimental designs, useful to tailor the system response for a specific need.

### Pulsatile shear and strain

The present microvessel-on-chip also allows for the application of pulsatile shear stress and strain, which better mimics the in vivo mechanical environment. Pulsatile strain requires rapid and reversible gel deformation. Because hydrogels are poroelastic materials, they are ideal candidates for pulsatile stimulus generation. To impose pulsatility, a sinusoidal inlet flow rate *Q*_*i*_ with the following waveform is imposed:

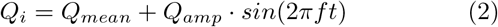

Setting both the mean flow rate *Q*_*mean*_ and the amplitude *Q*_*amp*_ to 10 µL min^−1^ and the frequency *f* to 1 Hz leads to significant oscillations of the channel diameter (Movie 4,5). The flow rate, wall shear stress, and channel strain all follow a sinusoidal waveform at the same frequency as *Q*_*i*_, with a small phase shift due to gel viscosity (Figure 7a, Movie 6). As expected, the cycle-averaged *Q*_*ch*_ oscillates; however, the amplitude of the oscillation is smaller than that of the imposed *Q*_*amp*_ (data not shown). While the amplitude of the oscillations in strain is virtually equal to that of the mean value, the oscillations in wall shear stress are considerably damped and are approximately half of the mean value (Figure 7a).

**Figure 7.**
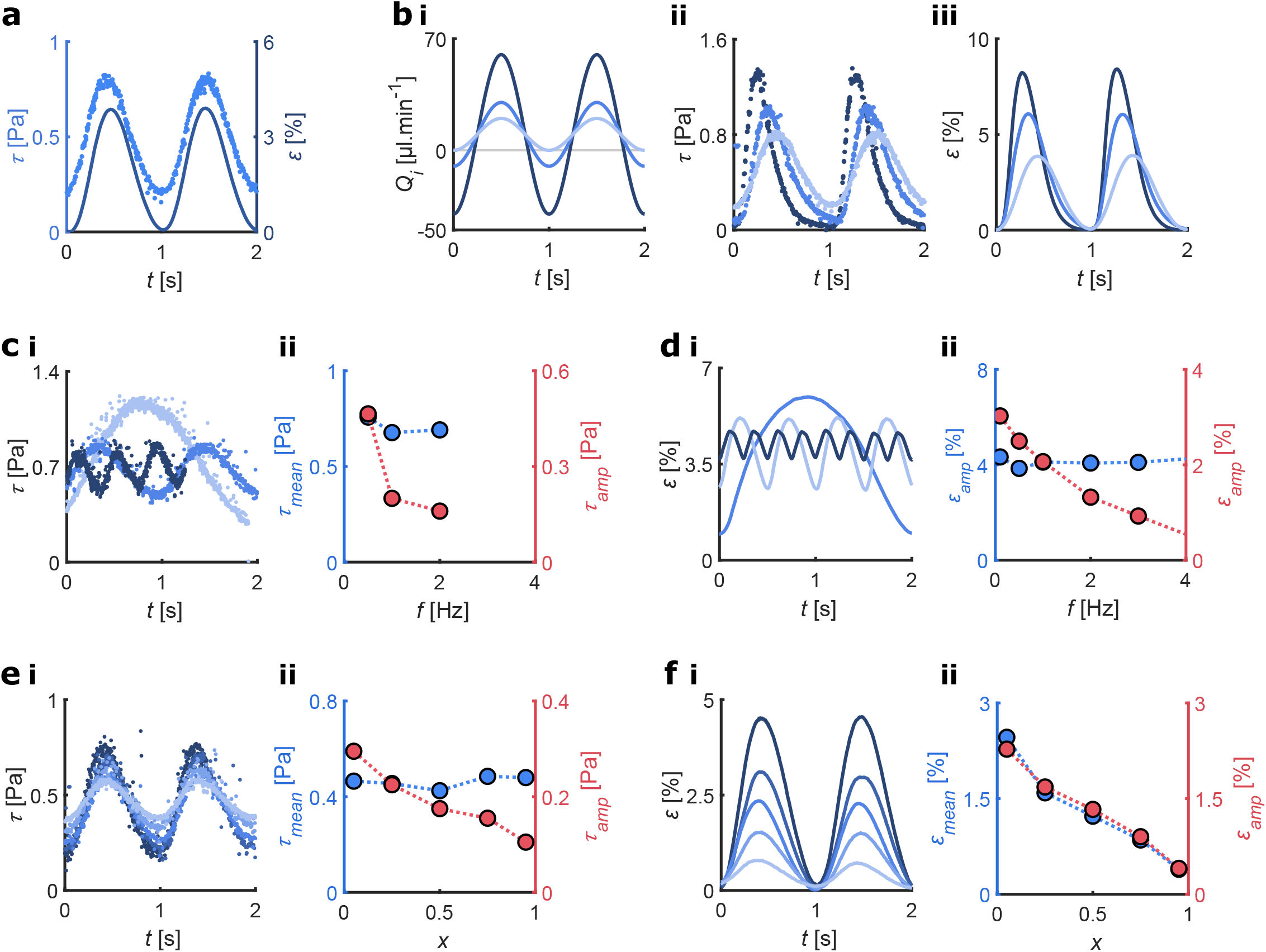
Response of the microvessel-on-chip to sinusoidal pulsatile flow. (**a**) Pulsatile stresses in the channel: mean wall shear stress *τ* (light blue, dots) and mean strain *ε* (dark blue, solid line) as a function of time for a sinusoidal imposed flow rate. (**b**) Impact of the amplitude of the imposed flow on channel response. Measured waveforms of the imposed flow rates (**i**), the resulting mean wall shear stress (**ii**), and mean strain (**iii**) (line colors match those of the corresponding imposed flow rate). (**c**) Wall shear stress waveform as a function of the sinusoidal frequency. **i** Measured waveforms for 0.5 Hz (light blue), 1 Hz (blue), and 2 Hz (dark blue). **ii** Measured mean (blue) and amplitude (red) of the wall shear stress oscillations as a function of frequency. (**d**) Strain waveform as a function of sinusoidal frequency. **i** Measured waveforms for 0.5 Hz (light blue), 2 Hz (blue), and 4 Hz (dark blue). **ii** Measured mean (blue) and amplitude (red) of the strain oscillations as a function of frequency. (**e**) Wall shear stress waveform as a function of position. **i** Measured waveforms from inlet (dark blue) to outlet (light blue). **ii** Measured mean (blue) and amplitude (red) of the wall shear stress oscillations as a function of position. (**f**) Strain waveform as a function of axial position. **i** Measured waveforms from inlet (dark blue) to outlet (light blue). **ii** Measured mean (blue) and amplitude (red) of the strain oscillations as a function of axial position. Dotted lines are guides for the eyes.

### Impact of the imposed waveform

To explore the extent of possible modulation of the amplitude of the pulsatile waveform for a given mean flow, *Q*_*amp*_ is set successively to 10, 20, and 50 µL min^−1^ while maintaining *Q*_*mean*_ at 10 µL min^−1^ and a 1 Hz frequency (Figure 7b.i). The amplitudes of the wall shear stress and the channel strain increase with increasing *Q*_*amp*_ as expected (Figure 7b.ii,iii). The mean wall shear stress increases slightly from 0.5 to 0.6 Pa and the mean strain increases from 2% to 4%, even though *Q*_*mean*_ is maintained constant (Figure 7b.ii,iii). Surprisingly, no negative values of wall shear stress or strain are observed. Additionally, the time behavior of both quantities deviates significantly from a sinusoidal waveform, with fast up and down phases followed by a short plateau phase at the end of the cycle. Tuning *Q*_*amp*_ allows control of the amplitude of the oscillations for both wall shear stress and strain. Interestinly, the cycle-average wall shear stress is less sensitive to *Q*_*amp*_ than the cycle-average strain.

### Impact of the imposed frequency

The frequency of the imposed flow rate controls the frequency of the response for both wall shear stress and strain. Shear stress and strain are recorded as functions of time for three physiologically relevant frequencies: 0.5, 1, and 2 Hz, with *Q*_*mean*_ and *Q*_*amp*_ held constant. The observed waveforms for both shear stress and strain remain sinusoidal, and the frequencies match those of the imposed signals (Figure 7c,d). The mean wall shear stress and strain are largely independent of frequency, while their amplitudes decrease with frequency (Figure 7c,d). Although the amplitudes of the oscillations decrease with frequency, significant amplitudes are obtained for the range of physiological frequencies studied, demonstrating the capabilities of the present system.

### Axial variations of the sinusoidal shear and strain

The mean value of pulsatile wall shear stress does not vary significantly along the channel length and remains around 0.46 Pa, whereas the amplitude of the shear stress oscillations decreases from 0.3 Pa to 0.1 Pa, characteristic of the progressive oscillation damping as described previously (Figure 7e). In contrast, both the mean and the amplitude of oscillation of strain decrease nearly linearly from 2.5% to 0.4% along the channel axis, with the amplitude equal to the mean value, as imposed through *Q*_*i*_ (Figure 7f). Strain results from compression of the hydrogel by the luminal pressure, and pressure exhibits a negative linear gradient from inlet to outlet for all time points. Therefore, the strain follows the same linear decrease along the channel length at each time point in accordance with the sinusoidal waveform.

## Discussion

In vivo, microvascular cells are subjected to both shear forces due to blood flow and stretch forces due to the transmural pressure difference. There is mounting evidence that when subjected simultaneously to multiple biophysical cues, cells integrate the information from the different cues and respond differently compared to cells subjected to one of the cues alone [57, 58]. Therefore, the ability to generate physiologically relevant shear and strain levels in the same system is particularly important. The microvessel-on-chip presented here accomplishes this objective via the novel approach of luminal flow actuation. While most existing microvessel-on-chip systems generate physiological shear stress levels, none are able to attain the strain levels reported here. Two unique features of our design, namely the dimensions of the hydrogel within which the microchannel is embedded and the open top of the chamber housing the gel, enable large strains. In current microvessel-on-chip systems, the thickness of the gel surrounding the channel is typically not more than a hundred microns, which greatly limits gel deformation. Here, the channel is positioned several hundred microns from all walls, thereby generating large strains in response to a relatively small increase in intraluminal pressure. Furthermore, most current systems are encased in closed chambers; therefore, an increase in intraluminal pressure translates into pressurization of the poroelastic hydrogel, where pore pressure and pore deformation are coupled, which resists deformation and limits strain. Opening the top of the hydrogel housing chamber is therefore essential for pressure release, thereby allowing large strains.

The experiments on the flow-actuated microvessel-on-chip are complemented with a simple analytical model based on an electrical circuit analogy combined with calculations of the hydraulic resistance of the porous media. The model employs an inverse problem approach whereby a comparison of the analytical results to the experimental data enables deducing otherwise unknown material properties of the system. Although the current model captures the principal experimental tendencies, incorporating additional complexity would further improve the predictive capabilities of the model. For instance, an equivalent mechanical circuit with a series of parallel springs, potentially non-linear, could be coupled to the equivalent electrical circuit to predict gel deformation. To this end, additional experimental data would be required to empirically determine the non-linear stress-strain relationship exhibited by the hydrogel. In the present model, the deformation is injected into the equations simply as a geometrical parameter.

Our experimental results indicate that the presence of an endothelium amplifies strain in the collagen hydrogel, an outcome explained by the barrier effect of the confluent monolayer to fluid transport. However, the cell monolayer is also an additional material with a certain stiffness, which would resist stretching. A more complete description of the mechanical contribution of the endothelium would therefore include its elasticity. Even though the dominant effect is the barrier effect, careful modeling of the non-linear poroelastic deformation of the gel would shed light onto a possible stiffening contribution of the ECs. Future modeling of the fluid-poroelastic structure interaction would also clarify the origin of the observed dynamic effects, most notably the modification of the sinusoidal waveform observed under pulsatile flow.

Previous in vitro platforms used to study the effect of strain on microvascular cells are poor mimics of the native environment, principally because they lack the coupling between strain and other biophysical factors such as flow-derived shear stress or substrate-derived cues including curvature, stiffness, and topography. The system presented here overcomes these limitations, enabling pulsatile shear and strain in a microvessel inside a soft fibrillar hydrogel. The extensive characterization presented demonstrates the complexity of the mechanical behavior of hydrogel-based in vitro microvessels and the inherent coupling that exists among the different mechanical stresses in these systems. The current study, which complements previous work on the use of hydraulic pressure for deforming hydrogels [59–61], establishes luminal flow actuation as a novel mechanism for generating physiologically and pathologically relevant strain levels in hydrogel-based microfluidic systems. The extensive investigation of different parameters provides a detailed operating manual that we hope will facilitate the adoption of flow actuation in future in vitro platforms.

In vivo, two additional mechanical stresses are present in the microvasculature: compressive stress due to the luminal pressure and shear stress on intercellular junctions due to transmural fluid flow. Both of these stresses have been shown to play important roles in EC mechanobiology [10, 58]. Generating these stresses requires porous medium flow through a hydrogel as well as control of the pressure gradient [62–64], features of which very few in vitro systems are capable. Although not detailed in this study, these two stresses are indeed present in the flow-actuated microvessel-on-chip presented here. The luminal pressure is around a few mmHg, corresponding to venous pressure levels. The transmural shear stress magnitude is difficult to evaluate as it depends on the exact geometry of cell-cell junctions [64]; it is nevertheless proportional to the flow loss, which we have characterized in detail here.

With the recognition of the critical importance of mechanical factors in regulating vascular structure and function, in vitro systems used in vascular biology research have evolved over the past two decades from simple flat petri dishes to microfluidic channels and most recently to vessel-on-chip systems where cells are subjected to controlled levels of shear stress. Accounting for wall deformation, however, requires more complex designs and has therefore remained largely elusive. We show that relatively simple design modifications of current microvessel-on-chip systems enable the use of luminal flow actuation as a novel strategy for attaining large wall deformations. The templating technique employed in the present work for producing microchannels in hydrogels is similar to that used elsewhere [46, 49] and has been applied to study the effects of flow on angiogenesis [47, 65], atherosclerosis [66], tumor vascularization and cancer cell dissemination [67,68], mural cell interactions [69], and blood-brain barrier function [70]. Adopting the flow actuation paradigm proposed here would enable the investigation of the role that wall strain plays in these important events and would thus add a new dimension to the ability of vessel-on-chip systems to mimic the in vivo mechanical environment.

## Conclusions

Although hydrogel-based microvessel-on-chip systems are increasingly popular mimics of the microvasculature, a major limitation of current systems is the inability to generate physiologically relevant levels of wall strain. In the present work, we show that combining specific design features with luminal flow actuation of a poroelastic collagen hydrogel constitutes a novel and highly effective strategy for overcoming these limitations. ECs lining the wall of this perfusable microvessel-on-chip can be subjected to a wide range of shear stresses and strains, steady or pulsatile, mimicking the native mechanical environment. Flow shear stress and wall strain in this system are strongly coupled, through the luminal pressure and the poroelastic nature of the hydrogel. We also show that a portion of the luminal flow seeps into the hydrogel, generating transmural flow that is associated with progressive luminal flow loss along the microvessel length. A key advantage of the flow actuation strategy described here is its simplicity and the fact that it obviates the need for external actuation.

Microvessel-on-chip systems have been shown to be excellent platforms for investigating the effect of flow-derived shear stress on a host of important phenomena such as endothelial-mural cell interactions, immune cell extravasation, tumor vascularization, and angiogenesis. Incorporating flow actuation into these systems provides the ability to also investigate the effects of wall circumferential stretch and thus greatly expands the mechanobiological toolbox. Beyond its application to the microvasculature, the concept of flow actuation of poroelastic hydrogels provides a general framework that promises to help unravel important open mechanobiological questions in many settings where cells are simultaneously subjected to mechanical stretching and interstitial shear.

## Acknowledgments

The authors thank Prof. Jean-Marc Allain for access to the OCT imager. The authors also acknowledge all members of the Barakat group for their constructive input during the project and the preparation of the manuscript. This work was funded in part by an endowment in Cardiovascular Bioengineering from the AXA Research Fund (to AIB) and an AMX doctoral fellowship from Ecole Polytechnique (to CAD).

## Contributions

**CAD** Conceptualization, Methodology, Investigation, Writing; **CRL** Investigation; **AB** Conceptualization, Methodology, Review & Editing; **AIB** Conceptualization, Review & Editing, Funding Acquisition.

## Appendices

### Appendix A: Model of the free and porous medium flow

As shown below, a system of two first order coupled differential equations is obtained by using an Ohm’s law analogy across *R*_*ch*_ and Kirchhoff’s current law at a node combined with Ohm’s law across *R*_*gel*_. *R*_*ch*_ is given by the hydraulic resistance for Poiseuille flow in a pipe and *R*_*gel*_ by Darcy’s law with a small modification to take the particular geometry of this system into account (see Appendix B). The two necessary boundary conditions are imposed by the experiment: the inlet flow rate is the imposed flow rate *Q*_*i*_ and the gauge pressure at the outlets is set to zero.

The hydraulic resistance *R*_*ch*_ for fully developed laminar flow in a cylindrical channel (Poiseuille flow) is given as:

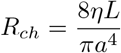

where *η* is the dynamic viscosity of the fluid, *L* the channel length, and *a* the channel radius.

As the channel deforms due to the luminal pressure and the pressure decreases axially, *a* is not constant. We assume a linear dependence on the axial position *x*: if *a*_*mean*_ denotes the average radius and assuming a 10% difference (based on the experimental measurement of strain, shown in Figure 3) between inlet and outlet, we express *a*(*x*) as:

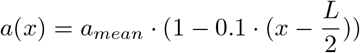

Consequently, the infinitesimal *R*_*ch*_, the resistance of a portion of the channel of length *dx* at position *x*, also depends on *x* and can be expressed as:

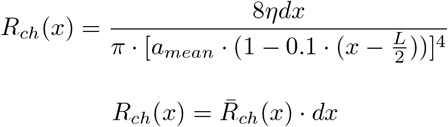

where

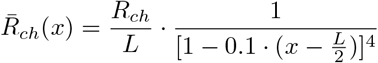

The flow resistance in the porous gel is given by Darcy’s law (see Appendix B). As the variations of channel diameter are small and the channel radius is an order of magnitude smaller than the gel dimensions, *R*_*gel*_ is assumed to be independent of x.

Applying Ohm’s law to one infinitesimal channel resistance (Figure 8a) yields:

**Figure 8.**
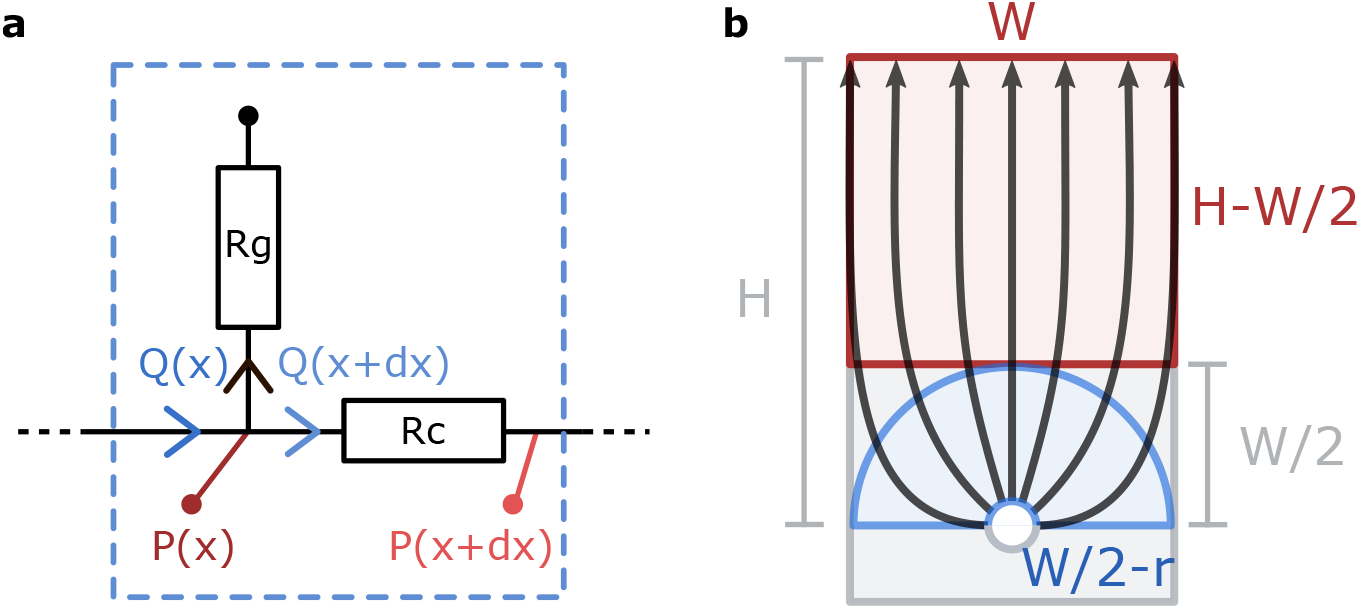
(**a**) Schematic of the equivalent electrical circuit. (**b**) Schematic of the equivalent geometries used to calculate the hydrogel hydraulic resistance showing the half torus in blue and the rectangle in red.

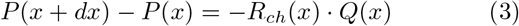

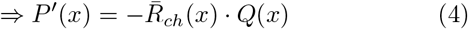

Using Kirchhoff’s current law at a node (Figure 8a), we obtain:

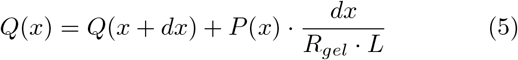

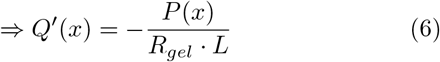

Combining equations (4) and (6) and the boundary conditions gives the following system of coupled differential equations:

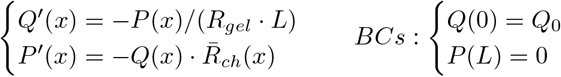

Normalizing the axial position by L leads to:

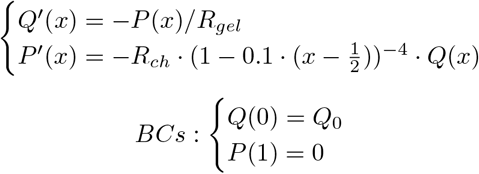

This system of coupled nonlinear differential equations is then solved numerically using the bvp4c Matlab solver.

To account for the cell layer in the case of an endothelialized channel, *R*_*gel*_ is replaced by *R*_*gel*_ + *R*_*cell*_, *R*_*cell*_ being the unknown resistance of the cell layer to be determined by fitting the experimental data.

To account for the additional outlet resistance *R*_*out*_, the boundary conditions are changed to the following:

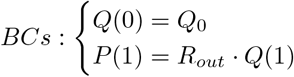

### Appendix B: Analytical expression for *R*_*gel*_

For simple geometries where diffusion occurs along only one dimension, Darcy’s law can be integrated to calculate the resistance of the porous medium in terms of the fluid viscosity *η*, the permeability *k*, and geometrical parameters. For diffusion across a rectangle of length *H*, width *W*, and thickness *L*:

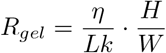

For diffusion across half a torus of inner radius *a*, outer radius *a*+*Δa*, and thickness *L*:

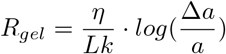

To derive an analytical expression, *R*_*gel*_ is approximated as two resistances in series: *R*_*gel1*_ models the initial radial diffusion through a torus and *R*_*gel2*_ models the final longitudinal diffusion through a rectangle (Figure 8b). As such, both resistances can be expressed as functions of the geometrical parameters of the system:

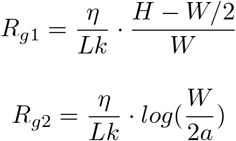

