## Supplementary Files for "Luminal Flow Actuation Generates Coupled Shear and Strain in a Microvessel-on-Chip"

### Supplementary figures

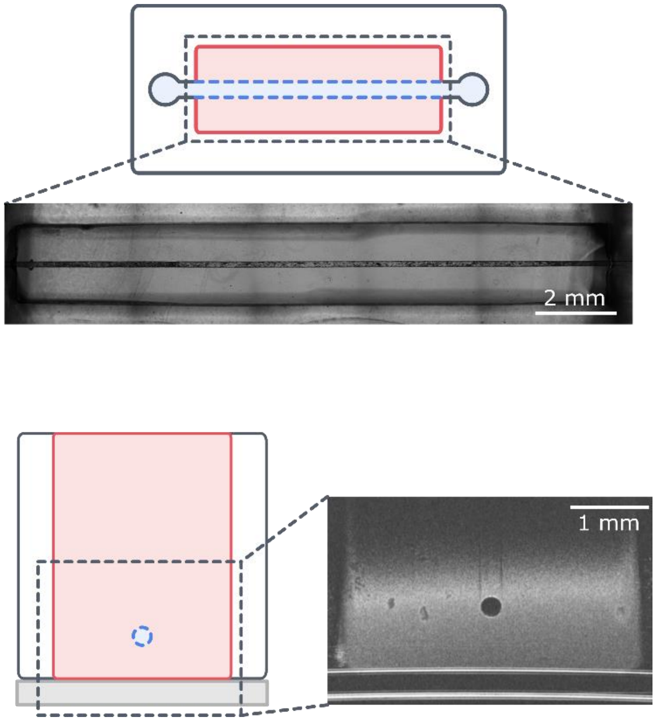


**Figure S1**. Illustrations and imaging of the entire microvessel-on-chip. (**a**) Top view of the microvessel-on-chip, brightfield imaging. (**b**) Cross-sectional view of the microvessel-on-chip, OCT imaging. The hydrogel is represented in red and the channel in blue.

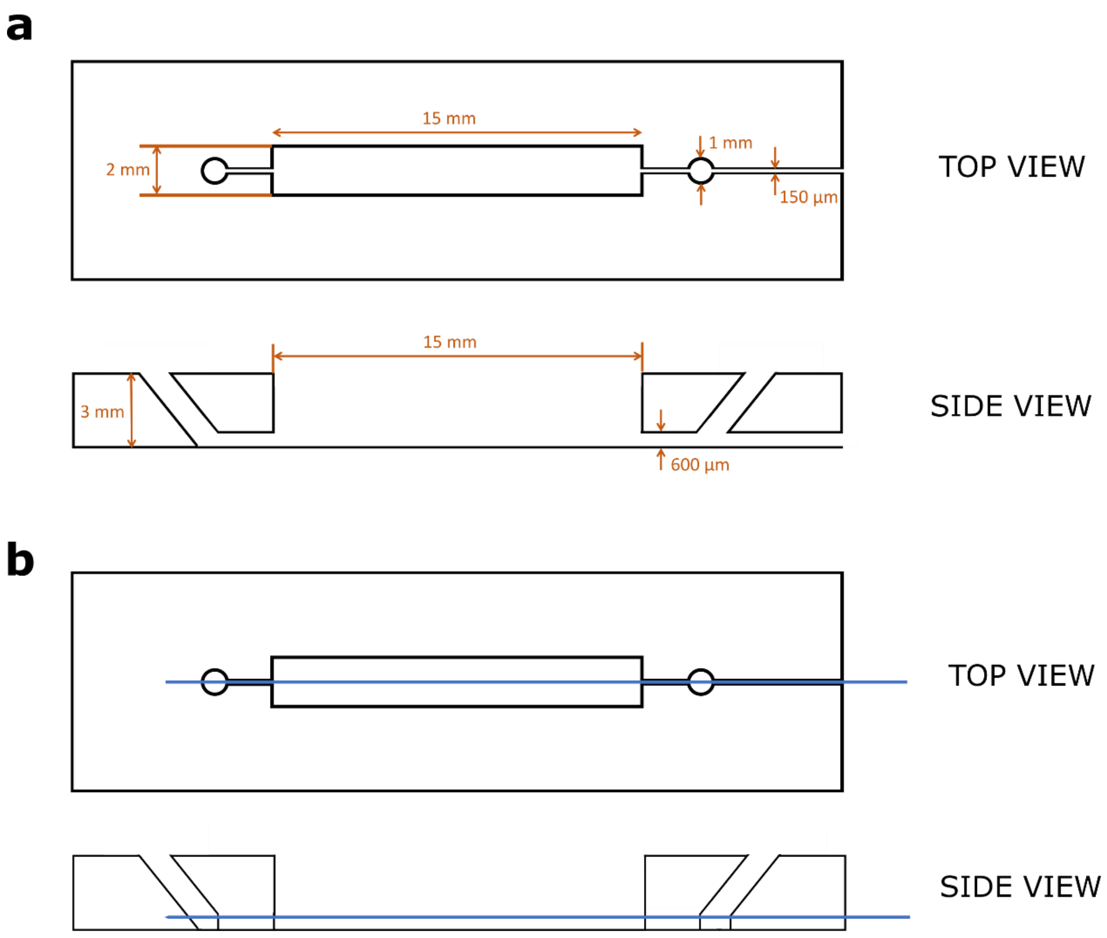


**Figure S2**. Schematics of the microvessel mold. (**a**) Geometry and dimensions of the PDMS mold. (**b**) Schematics of the needle (blue line) placement in the PDMS mold.

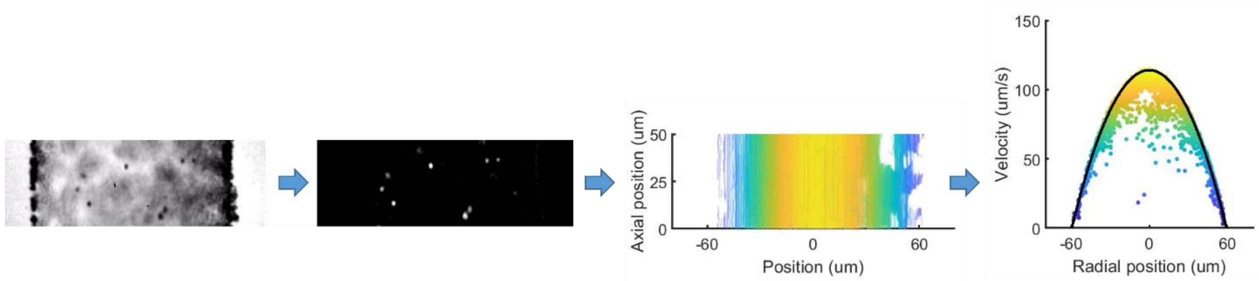


**Figure S3**. Particle tracking velocimetry process. The beads are first recorded with brightfield imaging followed by image processing and inversion. Individual tracks are then displayed as a function of radial position, and the mean velocity is extracted to reconstruct the parabolic profile.

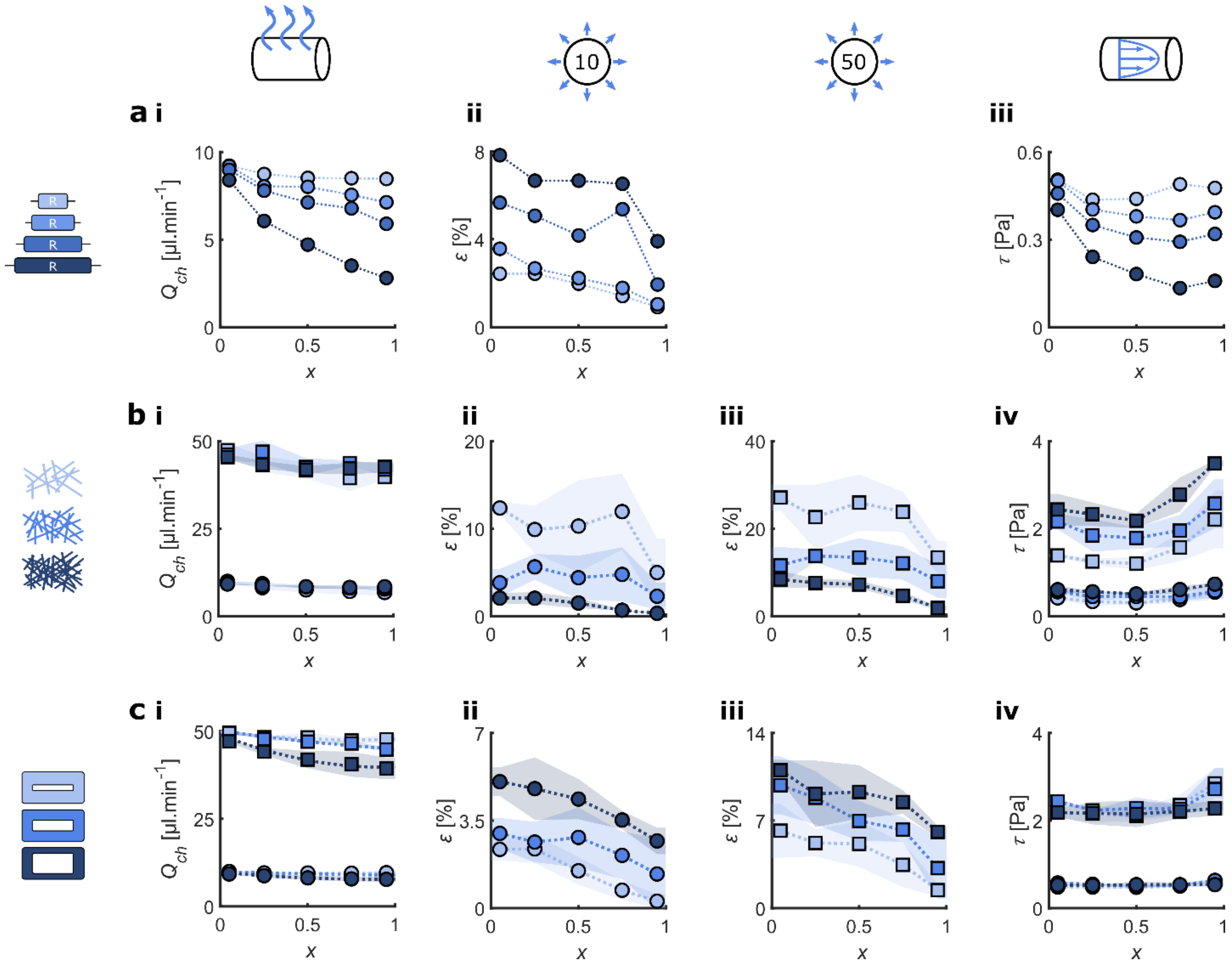


**Figure S4**. Axial variations for the three strategies used to modulate the shear-strain coupling: outlet resistance, gel width, and gel density. (**a**) Increasing the outlet resistance by increasing the length of the outlet tubing $L_{Rout}$. All results are for $Q_i = 10\mu L.min^{-1}$. **i** Flow in the channel demonstrating flow loss into the hydrogel. **ii** Strain $\varepsilon$. **iii** Wall shear stress $\tau$. (**b**) Changing the gel width. All results are repeated for the two imposed flow rates. **i** Flow in the channel demonstrating flow loss into the hydrogel. **ii** Strain $\varepsilon$ for $Q_i = 10\ \mu L.min^{-1}$. **iii** Strain $\varepsilon$ for $Q_i = 50\ \mu L.min^{-1}$. **iv** Wall shear stress $\tau$. (**c**) Changing the gel width. All results are repeated for the two imposed flow rates. **i** Flow in the channel demonstrating flow loss into the hydrogel. **ii** Strain $\varepsilon$ for $Q_i = 10\ \mu L.min^{-1}$. **iii** Strain $\varepsilon$ for $Q_i = 50\ \mu L.min^{-1}$. **iv** Wall shear stress $\tau$. All results are repeated for the two imposed flow rates. In all panels circles correspond to $Q_i = 10\ \mu L.min^{-1}$ and squares correspond $Q_i = 50\ \mu L.min^{-1}$. Dotted lines are guides for the eyes.

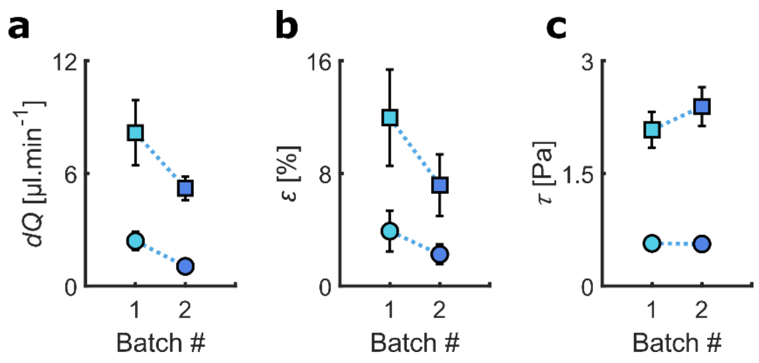


**Figure S5**. Influence of collagen quality on (**a**) flow loss *dQ*, (**b**) strain $\varepsilon$, and (**c**) wall shear stress $\tau$. In all panels circles correspond to $Q_i = 10\ \mu L.min^{-1}$ and squares correspond $Q_i = 50\ \mu L.min^{-1}$. Dotted lines are guides for the eyes.

### Supplementary movies

**Movie 1**. Animation of the PTV process for sinusoidal flow. Top left shows the live recording, bottom left the instantaneous velocity profiles, top right the flow rate as a function of time, and bottom right the 3D velocity profile.

**Movie 2**. Immunostaining of actin (phalloidin, white) and cell nuclei (DAPI, blue), demonstrating the confluent monolayer lining the microvessel lumen.

**Movie 3**. Microbeads suspended in PBS flown in the microchannel to visualize the flow.

**Movie 4**. Brightfield of the microchannel cross-section during a pressure step. Cells are visible on the edge of the channel as smooth bumps.

**Movie 5.** Brightfield and OCT imaging of the microchannel cross-section during sinusoidal oscillations at 1 Hz. Cells are visible on the edge of the channel in brightfield as smooth bumps.

**Movie 6.** OCT imaging of the microchannel cross-section during sinusoidal oscillations at 1 Hz. Cells are visible on the edge of the channel as the white lining.
